# The AIRE G228W mutation disturbs the interaction of AIRE with its partner molecule SIRT1

**DOI:** 10.1101/2021.08.24.457565

**Authors:** Jadson C. Santos, Mariangela Dametto, Ana Paula Masson, Vitor M. Faça, Rodrigo Bonacin, Eduardo A. Donadi, Geraldo Aleixo Passos

**Author notes:** Corresponding author at: Molecular Immunogenetics Group, Department of Genetics, Ribeirão Preto Medical School, University of São Paulo (USP), Ribeirão Preto, SP, Brazil. Equal contribution (Jadson C. Santos, Mariangela Dametto).

## Abstract

The autoimmune regulator (AIRE) protein functions as a tetramer, interacting with partner proteins to form the “AIRE complex,” which relieves RNA Pol II stalling in the chromatin of medullary thymic epithelial cells (mTECs). AIRE is the primary mTEC transcriptional controller, promoting the expression of a large set of peripheral tissue antigen genes implicated in the negative selection of self-reactive thymocytes. Under normal conditions, the SIRT1 protein temporarily interacts with AIRE and deacetylates K residues of the AIRE SAND domain. Once the AIRE SAND domain is deacetylated, the binding with SIRT1 is undone, allowing the AIRE complex to proceed downstream with the RNA Pol II to the elongation phase of transcription. Considering that the *in silico* and *in vitro* binding of the AIRE SAND domain with SIRT1 provides a powerful model system for studying the dominant SAND G228W mutation mechanism, which causes the autoimmune polyglandular syndrome-1, we integrated computational molecular modeling, docking, dynamics between the whole SAND domain with SIRT1, and surface plasmon resonance using a peptide harboring the 211 to 230 residues of the SAND domain, to compare the structure and energetics of binding/release between AIRE G228 (wild-type) and W228 (mutant) SAND domain to SIRT1. We observed that the G228W mutation in the SAND domain negatively influences the AIRE-SIRT1 interaction. The disturbed interaction might cause a disruption in the binding of the AIRE SAND domain with the SIRT1 catalytic site, impairing the AIRE complex to proceed downstream with RNA Pol II.

## Introduction

The human autoimmune regulator (*AIRE*) gene spans 14 exons at chromosome 21q22.3, encoding a 1635-nucleotide mRNA and a 545-amino acid protein (Aaltonen et al., 1994; Nagamine et al., 1997; Blechschmidt et al., 1999). Nearly 145 single-nucleotide mutations or large deletions have been observed along the *AIRE* gene (Wolff and Oftedal, 2019; Besnard et al., 2021; Human Gene Mutation Database available at https://www.hgmd.cf.ac.uk/ac/). Some mutations in the *AIRE* gene are related to autoimmune polyendocrinopathy-candidiasis ectodermal dystrophy (APECED) (Finnish-German APECED Consortium 1997; Nagamine et al., 1997), also known as autoimmune polyglandular syndrome type 1 (APS-1) (OMIM entry # 240300). Most APS-1 mutations are inherited in an autosomal recessive manner, except for a group of monoallelic mutations clustered within the zinc finger domain of the first plant homeodomain (PHD1) (Oftedal et al., 2015; Wolff and Oftedal, 2019) and a G228W mutation at the SAND domain, which follows a dominant inheritance pattern (Cetani et al., 2001; Ilmarinen et al., 2005; Su et al., 2008).

The 56-kDa AIRE protein is a transcriptional regulator in medullary thymic epithelial cells (Giraud et al., 2012; Passos et al., 2018; Perniola, 2018; Giraud and Peterson, 2019), exhibiting five domains and four LXXLL motifs. The caspase activation and recruitment (CARD) domain is essential for AIRE dimerization. The nuclear localization signal (NLS) domain presents a nuclear localization signal that mediates AIRE migration from the cytoplasm to the nucleus. The Sp100, AIRE-1, NucP41/75, and DEAF-1 (SAND) domains promote protein-protein interactions and chromatin binding, which play a role in the nuclear speckle localization of AIRE, and the PHD1 and PHD2 domains function as histone code readers, interacting and supporting the process of chromatin decondensation (Anderson et al., 2016; Kumar et al., 2001; Perniola et al., 2014; Giraud and Peterson, 2019). During its action in chromatin, AIRE interacts with several partner proteins forming the “AIRE complex,” in which SIRT1 is one of the most critical deacetylating K residues in the SAND domain, permitting the proper function of AIRE (Incani et al., 2014; Abramson et al., 2010; Abramson and Husebye, 2016).

In addition to chromatin binding, the SAND domain may also have a regulatory role in the homomultimerization of AIRE (Ramsey et al., 2002a; Halonen et al., 2004). The G228W SAND domain mutation was the first dominant mutation associated with the aggressiveness of APS-1 (Cetani et al., 2001; Ilmarinen et al., 2005; Su et al., 2008; Wolff and Oftedal, 2019); however, the reason for its dominant character and the molecular mechanism of APS-1 pathogenesis are still elusive.

Considering that i) the deacetylation of SAND domain K residues mediated by the partner SIRT1 molecule is essential for AIRE function in releasing RNA Pol II and allowing it to proceed with the elongation phase of transcription (Vaquero et al., 2004; Giraud et al., 2012, Saare et al., 2012, Incani et al., 2014, Chuprin et al., 2015, Bheda et al., 2016), and ii) the G228W mutation occurs at a position located at six amino acid residues from one of the acetylated K residues (K222), which is further deacetylated by SIRT1 (Saare et al., 2012), in this study, we hypothesized that the proximity of the G228W mutation to the K222 residue might interfere with the interaction between the SAND domain and the SIRT1 protein.

We integrated computational molecular modeling, docking, and dynamics between the whole AIRE SAND domain with the SIRT1 protein to test this hypothesis. Additionally, we evaluated the binding of a peptide harboring the 211 to 230 residues of the SAND domain with the SIRT1 protein to compare the binding/release between AIRE G228 (wild-type peptide) and W228 (mutant peptide) using surface plasmon resonance. The *in silico* analyses showed differences in the physical interaction between the AIRE SAND domain and the SIRT1 catalytic site for both wild-type and mutant domains. On the other hand, the surface plasmon resonance evaluation showed a stronger affinity (as seen through binding and dissociation) of the wild-type SAND domain peptide with the SIRT1 protein compared to a weaker association observed for the mutant peptide.

## Materials and Methods

### Protein structure homology modeling

We used the SWISS-MODEL workspace for protein modeling (http://swissmodel.expasy.org/workspace). The amino acid sequence in the FASTA format of the SIRT1 target protein was used as input. The SWISS-MODEL algorithms performed BLASTp and provided the best homology models based on global model quality estimation (GMQE) and qualitative model energy analysis (QMEAN) statistical parameters. The GMQE provided the quality by combining the target-template alignment properties, estimating a range between 0 and 1. The scoring function of QMEAN consisted of a linear combination of structural descriptors, which is related to the high-resolution X-ray crystal structure, estimating a *Z*-score scheme.

### Molecular docking

Molecular docking analysis of the modeled AIRE SAND domain, wild-type or G228W mutant AIRE SAND domain, and the SIRT1 protein (PDB code 4I5I) was performed to study the stability of the protein-protein interaction. The ClusPro 2.0 protein-protein docking server (Alekseenko et al., 2020) (available at https://cluspro.org/) was used. In the ClusPro server, the rigid body docking phase uses the PIPER docking program (https://www.schrodinger.com/products/piper), which relies on the fast Fourier transform (FFT) correlation approach. PIPER represents the interaction energy between two proteins using the formula *E* = *w1Erep+w2Eattr+w3Eelec+w4EDARS, w*here *E rep* and *E attr* denote the attractive and repulsive contributions to the van der Waals interaction energy, and *E elec* represents the electrostatic energy. *EDARS* is a pairwise structure-based potential; it primarily means desolvation contributions, i.e., free energy change by removing the water molecules from the interface. The coefficients *w 1, w 2, w 3,* and *w 4* define the molecular weights of the corresponding amino acid residues.

### Systems setup and molecular dynamics simulations

As mentioned above, the three-dimensional (3D) structures of the complexes formed with AIRE wild-type or with AIRE mutant peptide and SIRT1 generated in the docking section were used as initial configurations for the molecular dynamics simulations. In this work, calculations were performed to verify the stability and affinity of the complexes.

To mimic the interaction between the two peptides and SIRT1 protein, the K222 residue was acetylated using the parameters given in the AMBER Molecular Dynamics Package (https://ambermd.org/) for this residue. ACK and the connectivity library file for K222 residue were generated by tleap.

The two systems were solvated using the TIP3P water model and neutralized with four Na+ ions in the tleap tool implemented in AMBERTOOLS21 (https://ambermd.org/AmberTools.php). The whole system was truncated in a box with 15 Å between the edges of the periodic box and the molecules analyzed. An 8 Å cutoff particle mesh Ewald (PME) method was used to treat the long-range electrostatic interactions. AMBER molecular dynamics software and the ff19SB force field were used to prepare the systems and run the simulations.

Before running, the complexes were subjected to minimization in two stages using the steepest descent method followed by a conjugate gradient. In the first stage, the elements, peptides, and SIRT1 were fixed, and the position of the water molecules was minimized. After that, the whole system was minimized. A restrained thermalization phase with the microcanonical ensemble (NVE) was performed, increasing the temperature by 0.1 K until it reached 300 K using the Langevin thermostat and SHAKE algorithm (https://ambermd.org/) to constrain hydrogen bonds.

The production dynamics were carried out at constant pressure (NPT ensemble) with a time step of 2 fs. The total simulation time for each complex was 2700 ns, and frames were saved every 20 ps. The analyses were performed in the last 2650 ns after the systems were equilibrated. All calculations shown in the current work were performed using AMBERTOOLS21 scripts. The visualization programs used to create the figures of the molecules were VMD for LINUXAMD64, version 1.9.3, and PYMOL (https://pymol.org/2/).

### Surface plasmon resonance

The surface plasmon resonance (SPR) experiments were performed on Biacore Model T200 equipment (GE Healthcare Biosciences, Uppsala, Sweden). For the immobilization of human SIRT1 (catalog number S8846, Sigma–Aldrich, Saint Louis, MO), a CM5 sensor chip S series (GE Healthcare Biosciences) has carboxylic groups available for covalent coupling was used. All solutions used were purchased from the equipment supplier (GE Healthcare Biosciences). The procedures described here were performed automatically with the Biacore instrument using pre-established programs for binding amines in peptides or proteins.

Briefly, the carboxylic groups on the CM5 sensor chip surface were activated with EDC + NHS, forming the NHS ester reactive intermediate, which, after washing with HBS-N, readily reacted with amines dissolved in the immobilization solution. The human SIRT1 ligand was dissolved in 10 mM sodium acetate pH 4.0 plus 20 μM NAD+ (catalog number N1636, beta-nicotinamide adenine dinucleotide hydrate, Sigma–Aldrich), which was immobilized onto the Biacore sensor chip according to the manufacturer’s instructions.

To detect interactions between SIRT1 and AIRE WT or AIRE G228W mutant SAND domain peptides, different concentrations of peptides were injected onto the CM5 chip with immobilized SIRT1 protein. Peptide samples were injected sequentially from lowest to highest concentrations (20, 100, and 200 μM) with a contact time of 300 s, a 20 μl/min flow, and a dissociation time of 1200 s. The surface was regenerated between samples by injecting 2.0 M glycine pH 2.0 (30 s, 30 μl/min) followed by injection of 2.0 M NaCl (30 s, 30 μl/min). The stabilization period between injections after regeneration was 200 s.

The respective sequences of the twenty-amino acid peptide residues (Synbio Technologies, Monmouth Junction, NJ, USA) comprising positions 211 to 230 of the SAND domain of the human AIRE protein are as follows: from the C to N-terminus, AIRE wild-type (IQQVFESGGSK<u>K</u>CIQVGGEF) and AIRE G228W mutant (IQQVFESGGSK<u>K</u>CIQVGWEF). The underlined K residue represents the position where the acetyl radical was added. The experiment determined whether the K residue underlined was acetylated or not. The AIRE wild-type or AIRE G228W mutation data are available through GenBank NCBI (https://www.ncbi.nlm.nih.gov/) accession numbers (NM_000383.4): c.682G>T G [GGG] > W [TGG], coding sequence variant autoimmune regulator (NP_000374.1): p. Gly228TrpG (Gly) > W missense variant, DbSNP (https://www.ncbi.nlm.nih.gov/snp/rs121434257) and ClinVar (https://www.ncbi.nlm.nih.gov/clinvar/variation/3313/).

Kinetic data were analyzed with Anabel Analysis of Binding Events software (Krämer et al., 2019), available at (www.skscience.org/anabel). Data obtained with the different concentrations of AIRE WT or mutant peptides were combined in the same graph.

## Results

### Homology molecular modeling of the AIRE SAND domain

To obtain a separate 3D human AIRE SAND domain structure, we used the SwissModel tool to model proteins through homology. We used the SAND domain sequence of the human SP100 protein (PDB code 1h5p) for the modeling, which features 39.62% sequence identity to the human AIRE SAND domain. The SP100 protein SAND domain conserves the SKKCIQ amino acid sequence in the AIRE SAND domain, corresponding to the accurate interaction site between SAND and the SIRT1 protein. Thus, obtaining a 3D structure of the wild-type AIRE SAND domain and the AIRE SAND domain carrying the G228W mutation was possible. Considering that the G228W mutation is located in a loop region of the AIRE SAND domain (Figure 1), we may infer that this substitution does not cause local protein misfolding.

**Figure 1.**
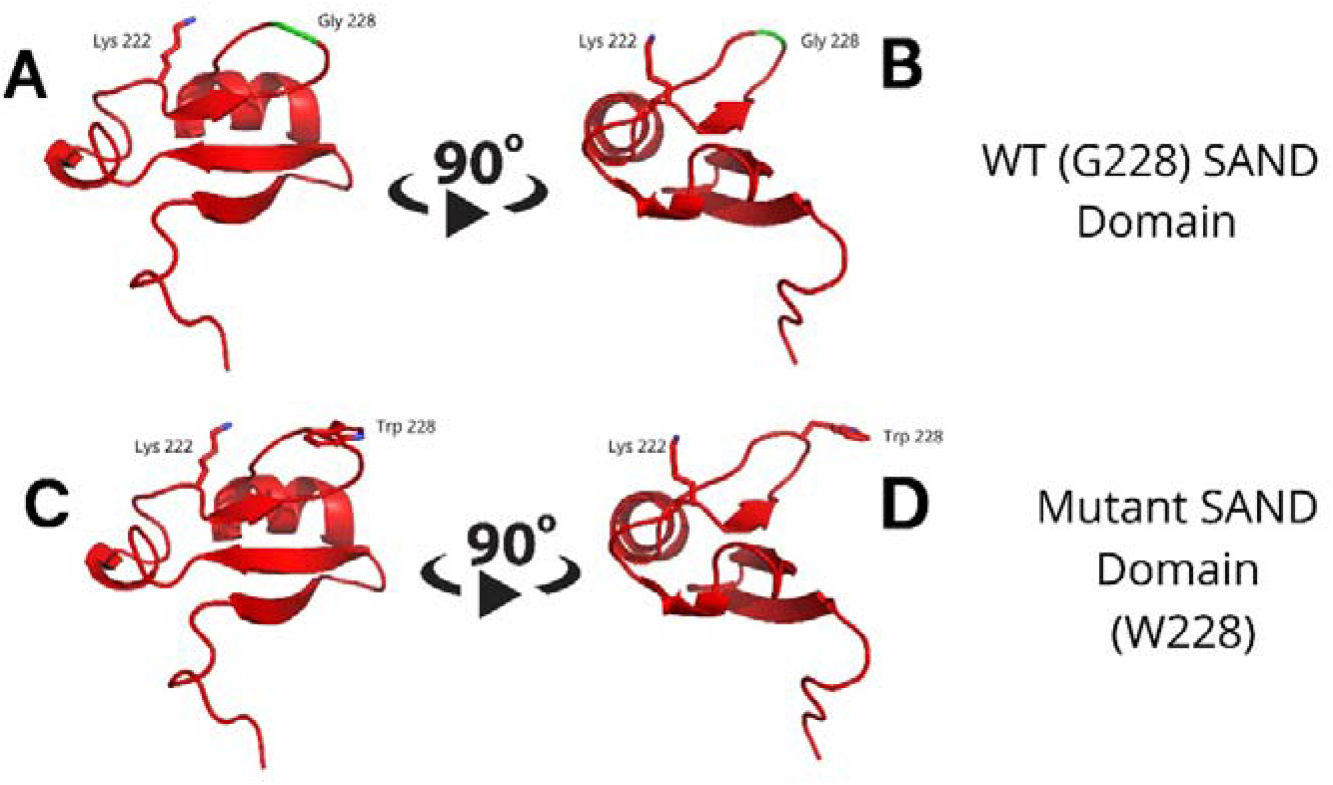
Molecular modeling of the G228 (Gly228) wild-type (A and B) or mutant W228 (Trp228) AIRE SAND domain (C and D) as acquired through homology modeling using the Swiss – Model with the human SP100 SAND domain as a template. The G228W mutation is located in a loop region and does not directly interfere with the formation of a secondary structure.

### Blind molecular docking shows that the SAND W228 mutant residue was targeted to the SIRT1 catalytic residue

To obtain an interaction model between the AIRE SAND domain (wild-type or G228W mutant) and SIRT1 and assess the affinity of the mutant SAND domain, we used the ClusPro 2.0 blind molecular docking tool. The docking was performed with nonacetylated K222 to understand whether the modeled AIRE SAND domain would have an affinity for the catalytic region of the SIRT1 protein.

Through blind molecular docking, it was possible to observe an affinity between the AIRE SAND domain WT G228 or W228 mutant domain and the catalytic region/site (H363 residue) of the SIRT1 protein, even without any previous targeting or chemical modification of the K222 residue, which is a target of the SIRT1 protein (Figure 2). Important to note that when K222 is nonacetylated, the structure formed is incompatible with the expected native structure.

**Figure 2.**
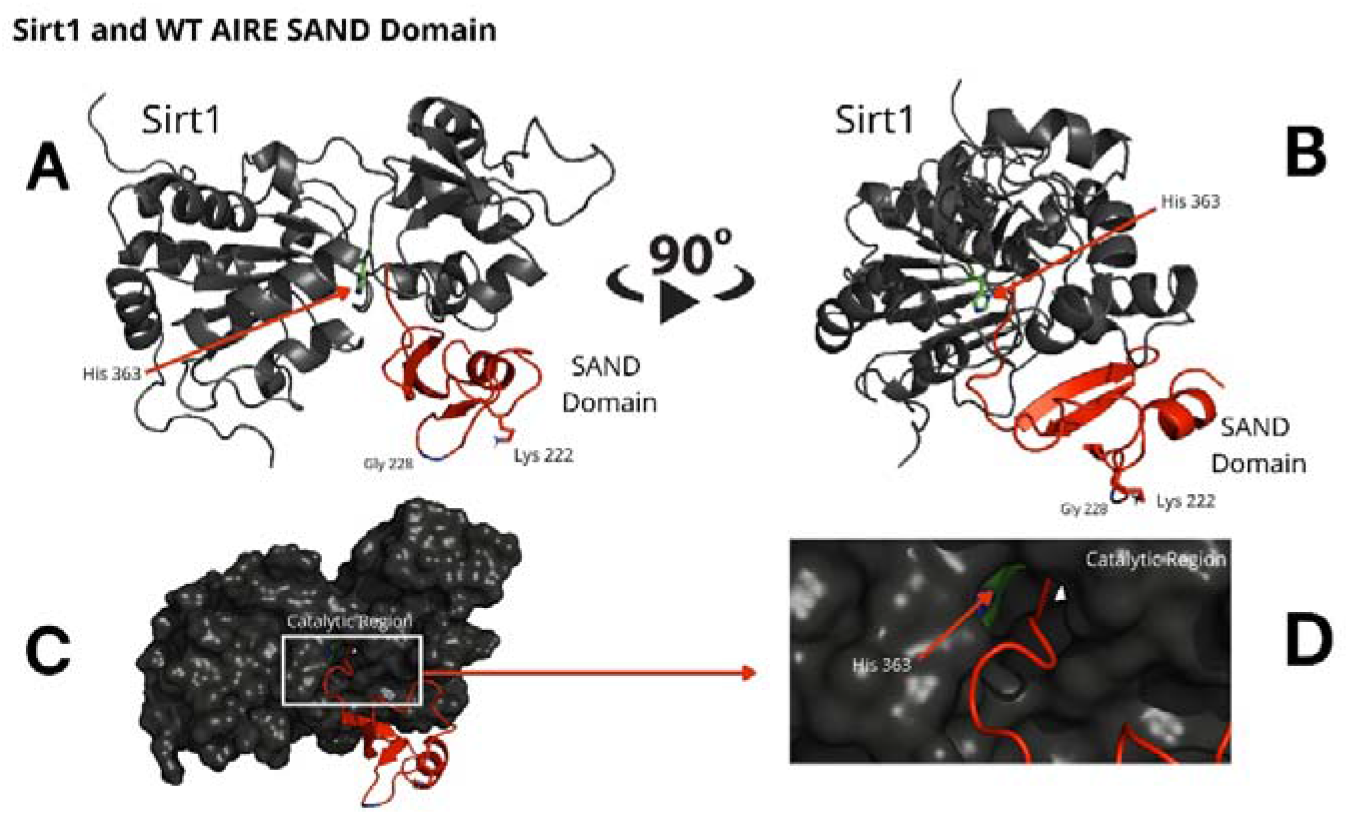
The blind docking output of the wild-type AIRE SAND domain shows that this domain interacts with the catalytic region of the SIRT1 protein. In A and B, the structures of the WT SAND domain (in red) and SIRT1 (in dark gray) are represented. In this representation, the AIRE SAND domain interacts with the catalytic region through its terminal loop, and the glycine 228 (Gly228) and lysine 222 (Lys222) residues are presented with their side chains. Without the Lys222 acetylated, this formation is incompatible with the native structure of the complex formed with the complete AIRE protein, in which this loop does not exist. In C and D, the same complex is represented with the surface of SIRT1 highlighted. D represents the catalytic region of SIRT1 and its catalytic residue, His363.

Comparing the structural conformations, we observed that the SAND domain mobilizes the W228 mutant residue that interacts with the SIRT1 H363 catalytic residue in the docking analyses (Figure 3).

**Figure 3.**
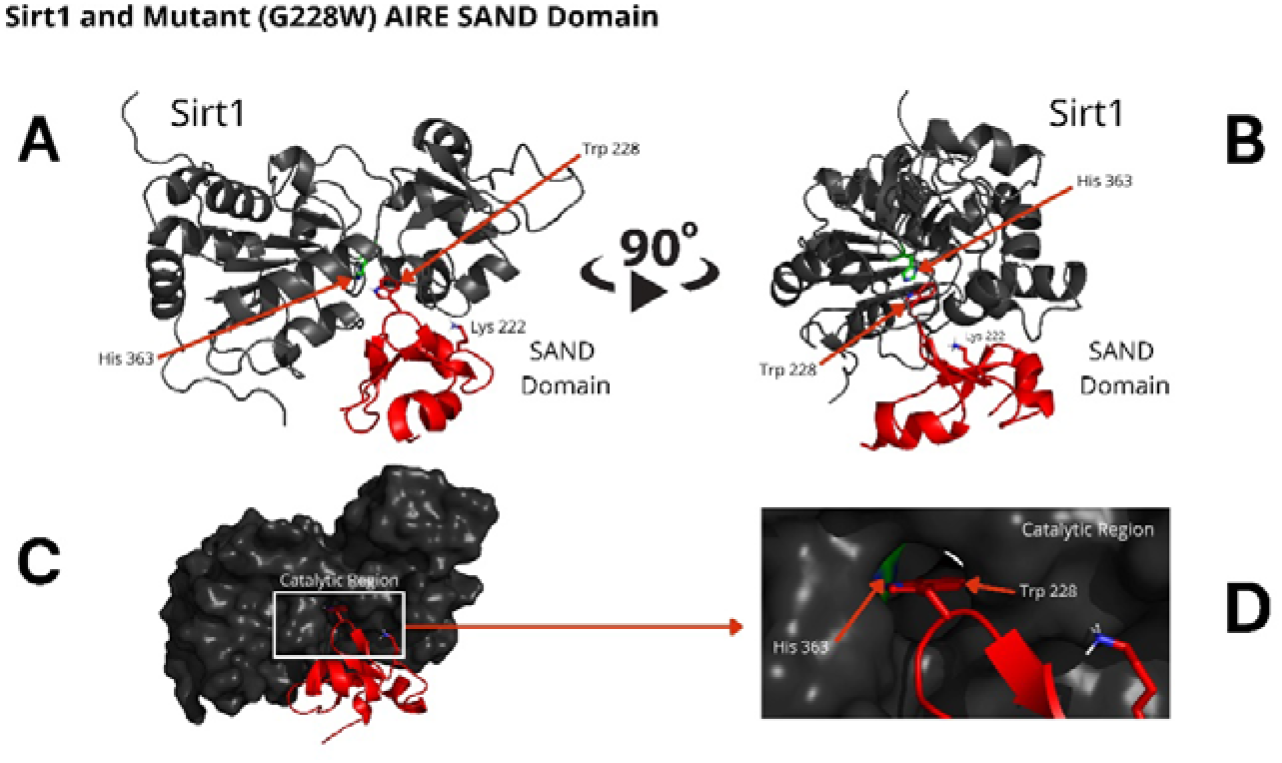
The blind docking output of the mutant AIRE SAND domain shows that this domain interacts with the catalytic region of the SIRT1 (His363 in green) protein through its mutant residue W228 (Trp228). The structures of the mutant SAND domain (in red) and SIRT1 (in dark gray) are represented in A and B. In this representation, the mutant AIRE SAND domain interacts with the catalytic region through its mutant residue W228 (Trp228), and the lysine 222 (Lys222) residues are presented with its side chain. In C and D, the same complex is represented with the surface of SIRT1 highlighted. D represents the catalytic region of SIRT1 and its catalytic residue, His363.

### Molecular dynamics

The molecular dynamics results with the AIRE SAND domain (wild-type or G228W mutant) showed that even the acetylated K222 residue at 1 μs of simulation did not allow the wild-type domain to form a stable structure, in which the K222 residue would interact with the H363 residue from the catalytic site of SIRT1. On the other hand, the G228W mutant domain exhibited a stable interaction with the SIRT1 catalytic site at the H363 residue. The analysis of the dynamics was performed to understand if K222 acetylation would be able to direct it to the catalytic residue of SIRT1.

The RMSD results showed that the WT SAND domain and SIRT1 complex is slightly less stable along the simulation time (1200 ns) than the complex formed between the SAND G228W mutant. Nevertheless, the G228W mutant domain complex remained stable in the RMSD [Figure 4, where higher angstrom distance values (Y-axis) indicate less stability]. The analysis of structures from molecular dynamics showed that even having inserted an acetyl radical at K222 residue in the WT or mutant domains, there was no targeting to the catalytic residue H363 of SIRT1 along the molecular dynamics simulation time. For molecular docking analyses, we used the SAND domain without chemical modification. Still, to understand the behavior of this domain as the SIRT1 protein recognizes it, we acetylated K222 for dynamic analysis (Figure 5).

**Figure 4.**
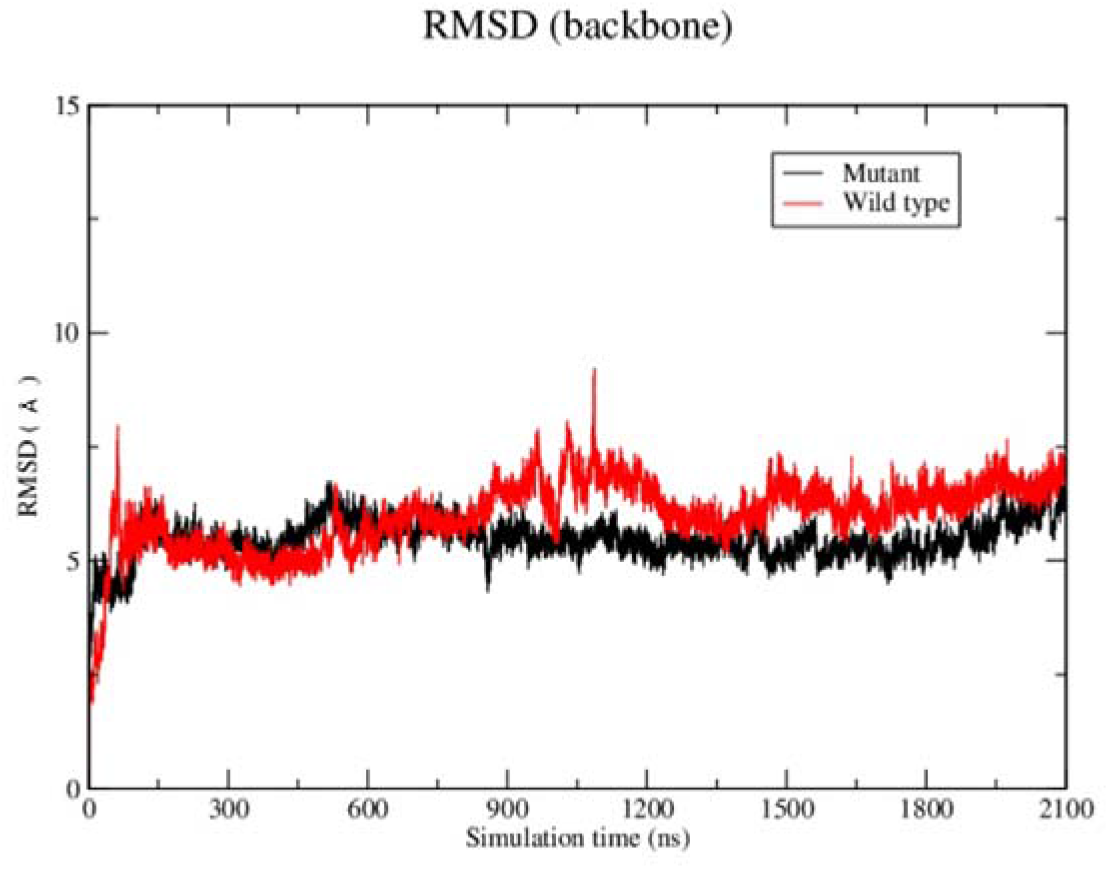
Root-mean square-deviation (RMSD) plot as a result of the molecular dynamics analysis of the wild-type (red) or mutant W228 (black) AIRE SAND domain with the SIRT1 protein. The RMSD plot shows that both WT and mutant SAND domains present a similar behavior along the first 600 ns. Then, the WT SAND domain exhibited a slight offset from 600 to 900 ns, and both complexes tended to stability at approximately 1200 ns.

**Figure 5.**
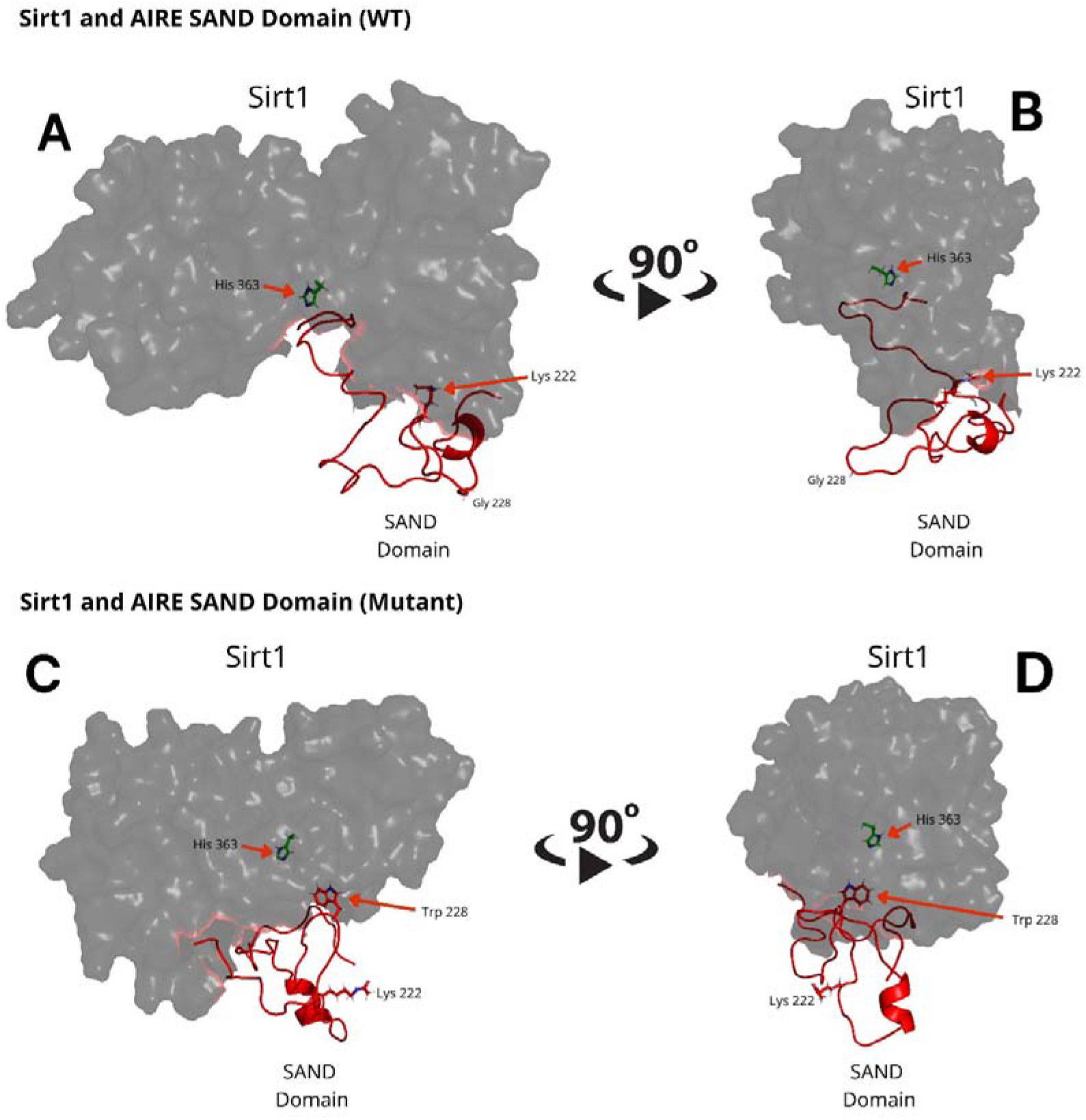
Molecular dynamics structures formed between (A and B) the WT AIRE SAND domain (in red) and SIRT1 protein (in dark grey) and (C and D) between Mutant AIRE SAND domain (in red) and SIRT1 (in dark grey). (A and B) Note that the acetylated K222 residue from the AIRE SAND domain (in red) is not directed to the SIRT1 H363 catalytic residue (side chain in green) after the analyzed dynamics time (1.0 microsecond) (C and D). Considering the complex formed between the mutant AIRE SAND domain, at 1.0 microseconds of dynamics, the mutant residue W228, in the AIRE SAND domain, remained close to the catalytic H363 residue (side chain in green) of SIRT1.

### Surface plasmon resonance indicates that the SAND G228W mutation modifies the interaction kinetics of AIRE with SIRT1

Considering that, the results from molecular docking and molecular dynamics exhibited slight differences when comparing WT and mutant SAND domains during the formation of complexes with SIRT1, next, we performed surface plasmon resonance (SPR) to study the behavior of the two types of peptides when they bind to SIRT1. The SPR results indicated a differential association and dissociation dynamics between the wild-type SAND G228 or mutant SAND W228 peptides and the SIRT1 protein. The interaction kinetics showed that the wild-type SAND G228 peptide has a more significant association capacity as observed at a concentration range of 20 to 200 μM. The interaction between the wild-type G228 SAND peptide and SIRT1 was almost completely dissociated after 1200 seconds of measurement. Conversely, the mutant W228 SAND peptide was less able to associate with SIRT1 (Figure 6). These results, combined with the *in silico* analyses, show that the G228W mutation changes the SAND-SIRT1 interaction, which is necessary to activate the AIRE protein. Of note, SPR results were in contrast with the blind docking and molecular dynamics, i.e., instead, the WT complex was more stable and mutant less stable. The SPR results were reproduced with six different concentrations of each of the acetylated peptides, WT and mutant (range 10 to 200 μM), in addition to the control (0 μM) (Supplementary Figure 1). For clarity of the results, we chose three concentrations (20, 100 and 200 μM) in addition to the control (0 μM), whose results we plot the graph in Figure 6.

**Figure 6.**
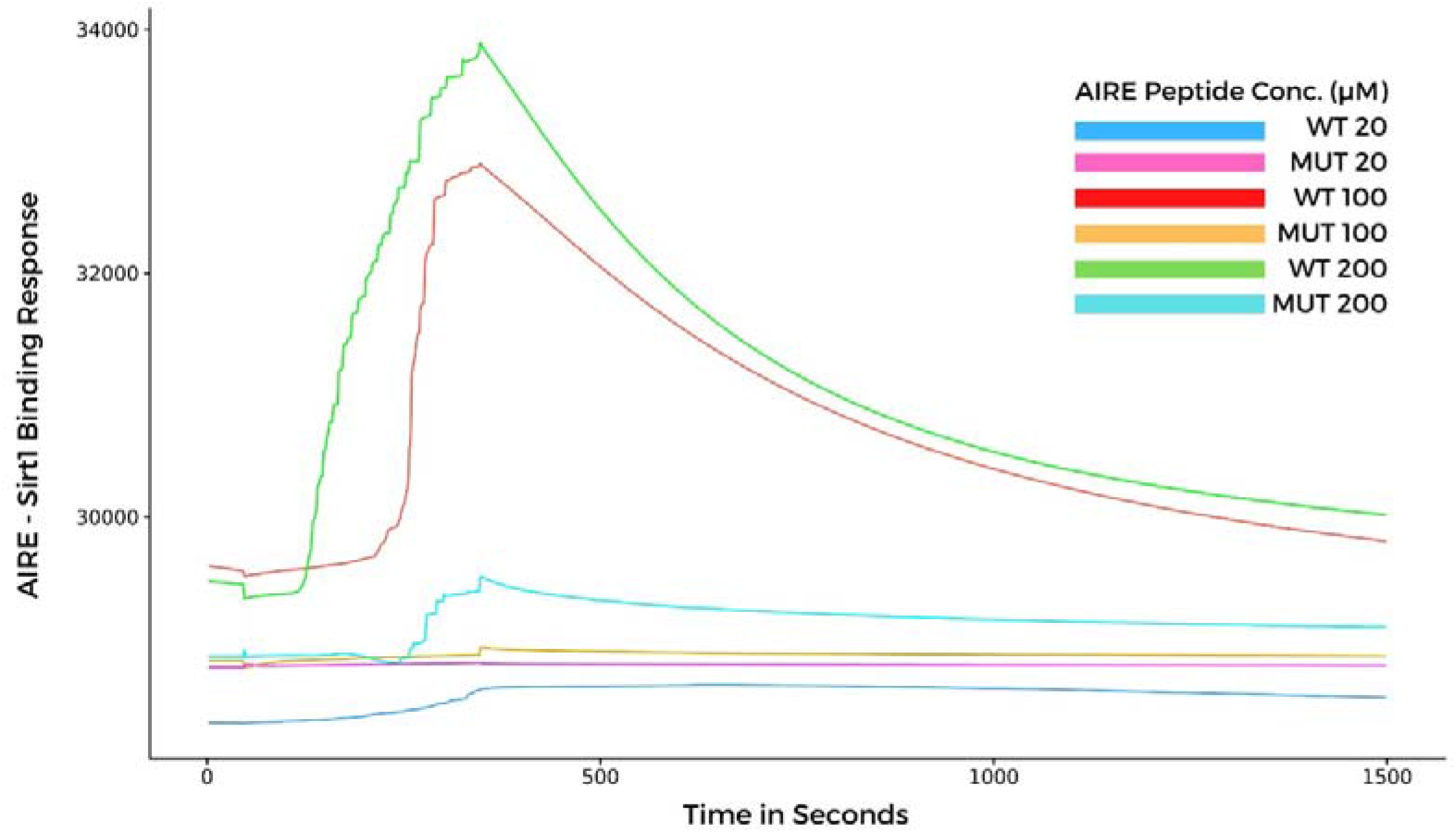
Surface plasmon resonance sensorgrams for binding the AIRE SAND domain with SIRT1 protein immobilized onto a Biacore sensor chip. The SAND domain peptides were assayed at various concentrations (20, 100, and 200 μM) with a contact time of 300 seconds, a 20 μl/min flow, and a dissociation time of 1200 seconds. Note that the WT AIRE SAND peptide (Gly228) associated and dissociated with SIRT1 throughout the analysis, while the mutant AIRE SAND peptide (Trp228) exhibited lesser association affinity with SIRT1 when compared to the WT peptide.

## Discussion

We reported that the dominant G228W mutation observed at the SAND domain of the human AIRE protein found in APS-1 patients changes interaction with the deacetylase SIRT1 compared to the AIRE wild-type protein. Because of the interference of the W228 mutant residue, the kinetics of the association and dissociation processes differ compared to the wild-type, which might explain the nuclear accumulation of the AIRE complex, as observed in AIRE mutant cells. Since the mutant AIRE protein does not properly stabilize in the AIRE complex, it may abnormally accumulate in the nucleus, as previously observed (Ilmarinen et al., 2005). The binding/dissociation kinetics between AIRE-SIRT1 is crucial, and any disturbance in these processes may destabilize the catalytic action of SIRT1 for AIRE protein deacetylation. Thus, the model system employed in this work improves the understanding of the molecular mechanism of the G228W mutation.

The study of the molecular mechanisms of primary immunodeficiencies associated with the development of autoimmune disorders has contributed to understanding the mechanisms of tolerance loss to self-antigens. Although recessive AIRE mutations at the PHD1 domain may occur at a ratio of 1:1000 in the general population (Anderson et al. 2014; Su et al. 2016), AIRE SAND G228W is the first dominant mutation associated with APS-1 (Cetani et al., 2001; Ilmarinen et al., 2005; Wolff and Oftedal, 2019). In the context of dominant inheritance, the central question is how these mutations influence the function of AIRE at the molecular level. Halonen et al. (2004) and Ilmarinen et al. (2005) observed that the G228W mutation quantitatively alters the nuclear location of the AIRE protein, impairing its transactivating property. Furthermore, Anderson and Su (2016) demonstrated that this mutation interferes with the localization of the AIRE protein, reducing the transcription of peripheral tissue antigens (PTAs).

This study evaluated whether the G228W mutation changes the interaction with SIRT1, an essential partner protein of the AIRE complex. We accessed the 3D structure for the AIRE SAND domain by homology modeling to the human SP100 protein SAND domain, which is available in a public database. We observed that the G228W replacement, which occurs in a loop region, might not result in misfolding of the AIRE SAND 3D structure since loop regions do not present secondary structures (Donate et al., 1996; Bottomley et al., 2001). The wild-type and the mutant AIRE SAND domain may not differ in their respective 3D structures, as performed by homology. This may be due to the segment’s characteristics in which the G228W mutation is present, taking the form of a flexible loop region (SAND amino acid residues 226-228). Therefore, the G228W mutation must involve another mechanism. Next, we asked whether the mutation could interfere with AIRE binding to one of its partner proteins. To initiate the downstream cascade of interactions, AIRE needs to be deacetylated, relieving RNA Pol II stalling in the chromatin. As SIRT-1 is the only deacetylase that integrates the AIRE complex, we next evaluated the binding of the AIRE SAND domain to SIRT1.

For this purpose, the structures of the wild-type SAND domain, the G228W mutant output from homology modeling, and the structure of the SIRT1 catalytic domain (PDB code 4I5I) were used. With the domains modeled, we used blind docking to understand how the complex between the SAND domains of AIRE WT and mutant would be formed with the SIRT1 protein.

We observed that the SAND domain mobilizes the W228 mutant residue that interacts with the SIRT1 H363 catalytic residue, changing with the interaction kinetics between these proteins, followed by SPR measurements. As observed in this study, due to the SAND G228W mutation, the AIRE protein might bind weakly to SIRT1. The AIRE complex, including RNA Pol II, formed with an AIRE mutant protein, might be dysfunctional and keep stalled to chromatin. The dysfunctional AIRE complex might occur due to the non-deacetylation of K222 residue, which is essential for the proper function of the AIRE protein. In addition, to signify a non-physiological situation, this finding might explain the accumulation of the AIRE complex in the nucleus of mutant cells, as previously reported (Halonen et al. 2004; Ilmarinen et al. 2005).

The AIRE protein plays a role in the cell nucleus as a tetramer (Abramson et al., 2010; Incani et al., 2014), in which the SAND domain of each subunit has K residues SIRT1 deacetylates. SIRT1 exerts its enzymatic activity, which depends on the amino acid residues near the acetylated K residues in the AIRE SAND domain (Garske et al., 2006). Accordingly, we hypothesized that the G228W mutation close to acetylated K residues could interfere with the SIRT1 interaction since it is an enzyme necessary to convert the acetylated to the nonacetylated and functional form of AIRE.

Although there is a prediction that the K222 residue is only moderately acetylated (Chuprin et al., 2015), previous work (Saare et al., 2012) showed that this amino acid residue is acetylated in the presence of the CBP and p300 II proteins, using mass spectrometry. Several lines of evidence stress the importance of K deacetylation in the proper interaction of AIRE with SIRT1: i) functional analyses of the G228W AIRE SAND mutation show that the subunit-containing acetylated K residue could form a hybrid tetramer unable to promote AIRE transactivation activity (Halonen et al. 2004; Ilmarinen et al. 2005), ii) AIRE mutations mimicking acetylated K243 or K253 residues in the SAND domain also reduced the transactivation activity and accumulated as nuclear bodies, whereas mutations mimicking nonacetylated K residue were functionally similar to wild-type AIRE (Saare et al. 2012), and iii) the AIRE SAND R247C mutation retains the ability to form complexes with wild-type AIRE and localize into the nucleus; however, it reduces the activation of downstream transcription through dominant-negative inhibition (Abbott et al. 2018). Notably, the R247C mutation occurs close to other K residues (K243 and K253) in the SAND domain that SIRT1 can deacetylate, also harming the transactivation function of AIRE.

This approach confirmed the hypothesis raised in this study, in which we consider that the AIRE G228W mutation changes the interaction with SIRT1, mainly when evaluated using SPR analysis. Sensor-based SPR is a label-free, sensitive, precise, and *in vitro* tool used for determining binding kinetics between biological molecules, including antigen-antibody or protein-protein interactions, among other molecular interactions (Nguyen et al., 2015; Wang et al., 2019; Soltermann et al., 2021; Ribeiro et al., 2021).

The complementary approaches used in this study (*in silico* and SPR) allowed us to evaluate the structure, energetic affinity, and association/dissociation kinetics of the complete AIRE SAND domain and peptides (wild-type and G228W mutation) with the SIRT1 protein. We observed that: i) both complete SAND domain can bind to the SIRT1 catalytic domain, ii) the binding kinetics of the G228W mutant peptide exhibited a weaker association when compared with its wild-type counterpart, and as a result, the wild-type SAND domain peptide bound faster and dissociated longer, and iii) taken together, these approaches provided evidence that the G228W mutation acts as a destabilizer for AIRE’s interaction with its partner SIRT1 and offers a more accurate molecular basis for clarifying how the mutant G228W AIRE protein loses its function.

## Acknowledgments

This work was funded by the following Brazilian research support agencies; Fundação de Amparo à Pesquisa do Estado de São Paulo (FAPESP, grant # 17/10780-4 to GAP and EAD), Conselho Nacional de Desenvolvimento Científico e Tecnológico (CNPq, grants # 305787/2017-9 and # 311304/2021-4 to GAP, and # 302060/2019-7 to EAD), and Coordenação de Aperfeiçoamento de Pessoal de Nível Superior (CAPES, Finance Code 001).

## Contributions

JCS: raised hypothesis, discussed the analysis tools, performed molecular docking, analyzed data, interpreted the results, and wrote the manuscript.

MD: discussed the analysis tools, performed molecular dynamics, analyzed data and interpreted the results.

APM: performed protein binding kinetics (SPR), and interpreted the results.

VMF: performed protein binding kinetics (SPR), and interpreted the results.

RB: performed molecular dynamics, analyzed data.

EAD: raised hypothesis, interpreted the results and wrote the manuscript.

GAP: conceived the study, raised hypothesis, organized the study pipeline, interpreted the results and wrote the manuscript.

---

**Supplementary Figure 1.**
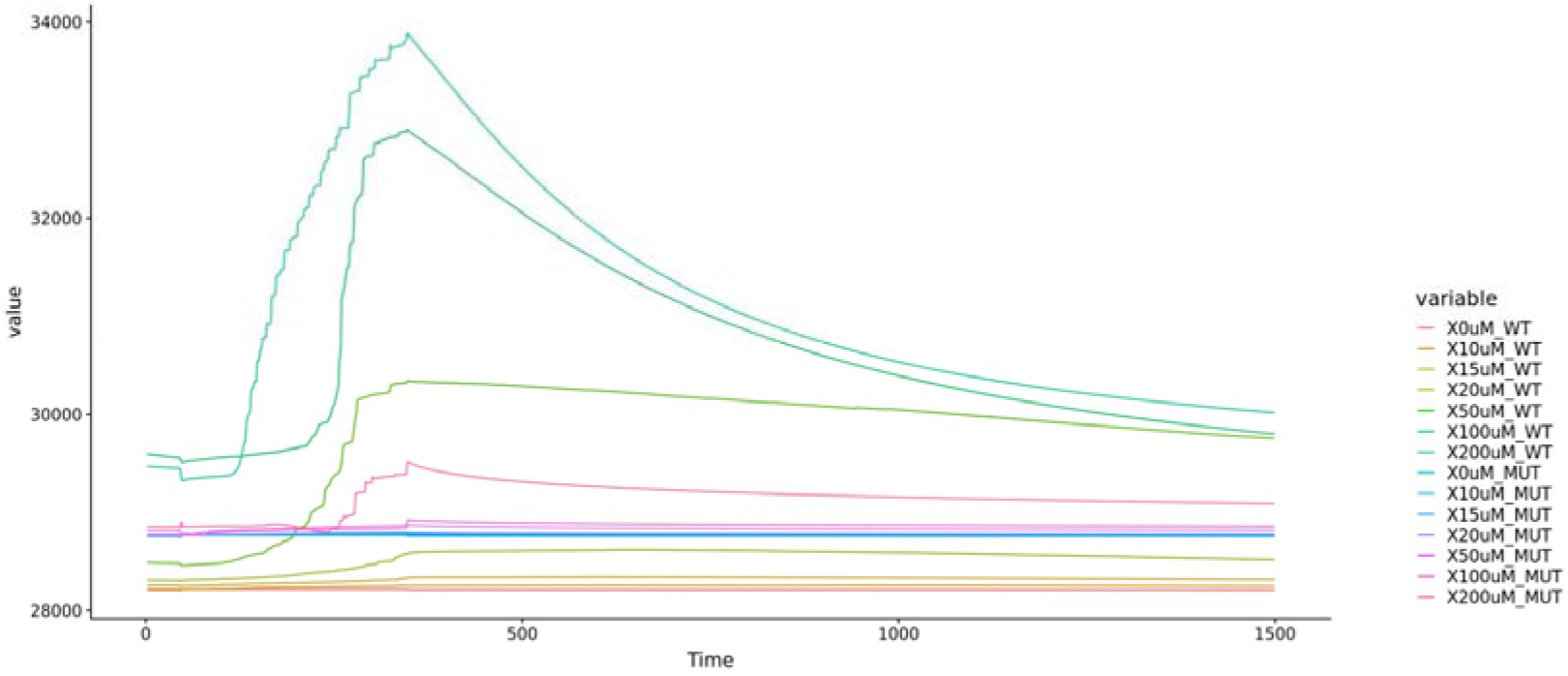
Surface plasmon resonance sensorgram analysis of the binding between WT or G228W mutant AIRE SAND domain peptides with SIRT1 protein immobilized onto a Biacore sensor chip. Six different peptide concentrations were assayed (range 10, 15, 20, 50, 100, and 200 μM) plus the control (0 μM) for both WT and mutant peptides. Binding analysis shows a progressive increase in the association with the WT peptide, which was not observed with the mutant peptide.

